# Paired single-cell imaging of calcium and expression to map niches of identity and function

**DOI:** 10.64898/2026.02.09.704922

**Authors:** Alexander P. Clark, Payton Gergen, Jeffrey J. Saucerman

## Abstract

Cellular identity is often inferred from molecular markers, while function is measured independently, obscuring how these dimensions align at single-cell resolution. In cardiomyocytes, this disconnect is especially limiting, as calcium dynamics and subtype markers are typically assessed in bulk or separate cells then averaged across populations. In human induced pluripotent stem cell–derived cardiomyocytes (iPSC-CMs), this gap has limited our ability to determine whether heterogeneity in electrophysiology and calcium handling reflects noise, maturation, or structured biological states. This lack of clarity is due in part to the lack of methods that directly link live functional measurements with molecular identity at single-cell resolution. Here, we introduce CARBONITE (<u>Ca</u>lcium <u>R</u>ecordings <u>B</u>ef<u>o</u>re Ide<u>n</u>tification by <u>E</u>xpression), a scalable single-cell imaging framework that pairs live calcium dynamics with protein expression and spatial phenotyping. Using high-content imaging in 96-well plates, CARBONITE integrates per-cell calcium transient features with immunofluorescent identification applied to cardiomyocyte subtype markers, enabling quantitative functional–molecular mapping within the same cells. Applying CARBONITE to mixed human induced pluripotent stem cell derived cardiomyocyte (iPSC-CM) populations reveals heterogeneity in calcium transient dynamics and marker expression within individual wells. Canonical atrial and ventricular markers capture only a subset of functional variability. Notably, calcium transient shape segregates cells into two discrete functional states that exhibit perinuclear ANP (atrial marker) enrichment and nucleation state (i.e., mono- vs. binucleation). Notably, these groups show know relationship to MYL2 (ventricular marker) expression. Binucleated cells are more likely to exhibit a spike-like calcium transient, identifying nucleation as a dominant and previously underappreciated axis of cardiomyocyte identity influencing calcium function. Together, these results establish CARBONITE as a functional multimodal single-cell platform that reveals organizational principles of cell identity, providing a foundation for dissecting functional niches in development and disease.

## Introduction

Cellular identity is commonly inferred from molecular markers, while functional behaviors are measured independently. This separation has enabled powerful reductions of complex biological systems, but it obscures how molecular state and dynamic function align at single-cell resolution. As a result, heterogeneity observed in functional assays is often difficult to interpret: it may reflect experimental noise, differences in maturation, or discrete and biologically meaningful states that are not captured by canonical markers.

This challenge is particularly acute in excitable cell types, where dynamic function is a defining feature of identity. Cardiomyocyte function arises from tightly regulated interactions between calcium handling, contractile machinery, and gene regulatory programs that define cellular identity. Human induced pluripotent stem cell–derived cardiomyocytes (iPSC-CMs) are widely used to study these relationships in the context of disease and pharmacology [1–5]. But, iPSC-CM cultures exhibit substantial cell-to-cell heterogeneity in electrophysiology and gene expression, even within nominally homogeneous wells [6–9]. The extent to which this heterogeneity reflects experimental artifact, maturation differences, or discrete biological states remains poorly understood [7,10,11].

Calcium transients are typically analyzed at the field-of-view or well level, obscuring single-cell function [12,13]. At the same time, chamber-specific (e.g., atrial vs. ventricular) cardiomyocyte identity is commonly inferred using canonical markers such as perinuclear atrial natriuretic peptide (ANP) and ventricular myosin light chain 2 (MYL2) expression or action potential (AP) shape [7,14,15]. These marker- and AP-based classifications are often assumed to correspond, yet this assumption has rarely been tested at single-cell resolution. As a result, functional and molecular phenotyping are frequently treated as coupled, despite being measured independently [13].

Single-cell RNA-seq and ATAC-seq have demonstrated that cardiomyocyte identity is multidimensional, shaped by intersecting transcriptional and epigenetic axes. These approaches have transformed our understanding of cellular heterogeneity by revealing both concordant and discordant features across molecular modalities [16]. By necessity, such measurements decouple molecular state from live functional behavior. Conversely, live-cell calcium imaging provides rich, quantitative descriptions of dynamic cardiomyocyte function but lacks paired molecular information within the same cells. This disconnect has limited efforts to directly map functional states to molecular identity and local microenvironment.

Here, we introduce CARBONITE (<u>Ca</u>lcium <u>R</u>ecordings <u>B</u>ef<u>o</u>re Ide<u>n</u>tification by <u>E</u>xpression), a scalable single-cell imaging framework that pairs live calcium dynamics with protein expression and spatial phenotyping in intact iPSC-CM cultures. Conceptually analogous to multimodal single-cell genomics, CARBONITE integrates quantitative functional, molecular, and spatial measurements at single-cell resolution. Applying this approach to mixed human iPSC-CM populations, we uncover substantial heterogeneity in calcium transient shape and demonstrate that canonical atrial and ventricular markers are differentially—but incompletely—associated with functional state. Notably, we identify a dominant axis of calcium transient organization linked to ANP enrichment and nucleation status (mono- vs. multinucleation), revealing an unexpected dimension of cardiomyocyte identity not correlated to MYL2 expression.

Together, these results establish CARBONITE as a generalizable platform for resolving functional niches in cardiomyocyte populations and underscore the importance of paired single-cell functional–molecular measurements for understanding cardiac cell identity.

## Results

### CARBONITE is a quantitative single-cell imaging framework for paired calcium and protein expression

To enable linkage between cardiomyocyte functional dynamics and molecular identity at single-cell resolution, we developed **CARBONITE** (Calcium Recordings Before Identification by Expression), a quantitative imaging framework that pairs live-cell calcium recordings with protein expression measurements in the same cells (**Figure 1**). CARBONITE was designed to capture spontaneous calcium dynamics in iPSC-CMs prior to fixation, followed by immunofluorescent identification of cardiomyocyte subtype markers, enabling joint analysis of functional and identity-related features within individual cells.

**Figure 1:**
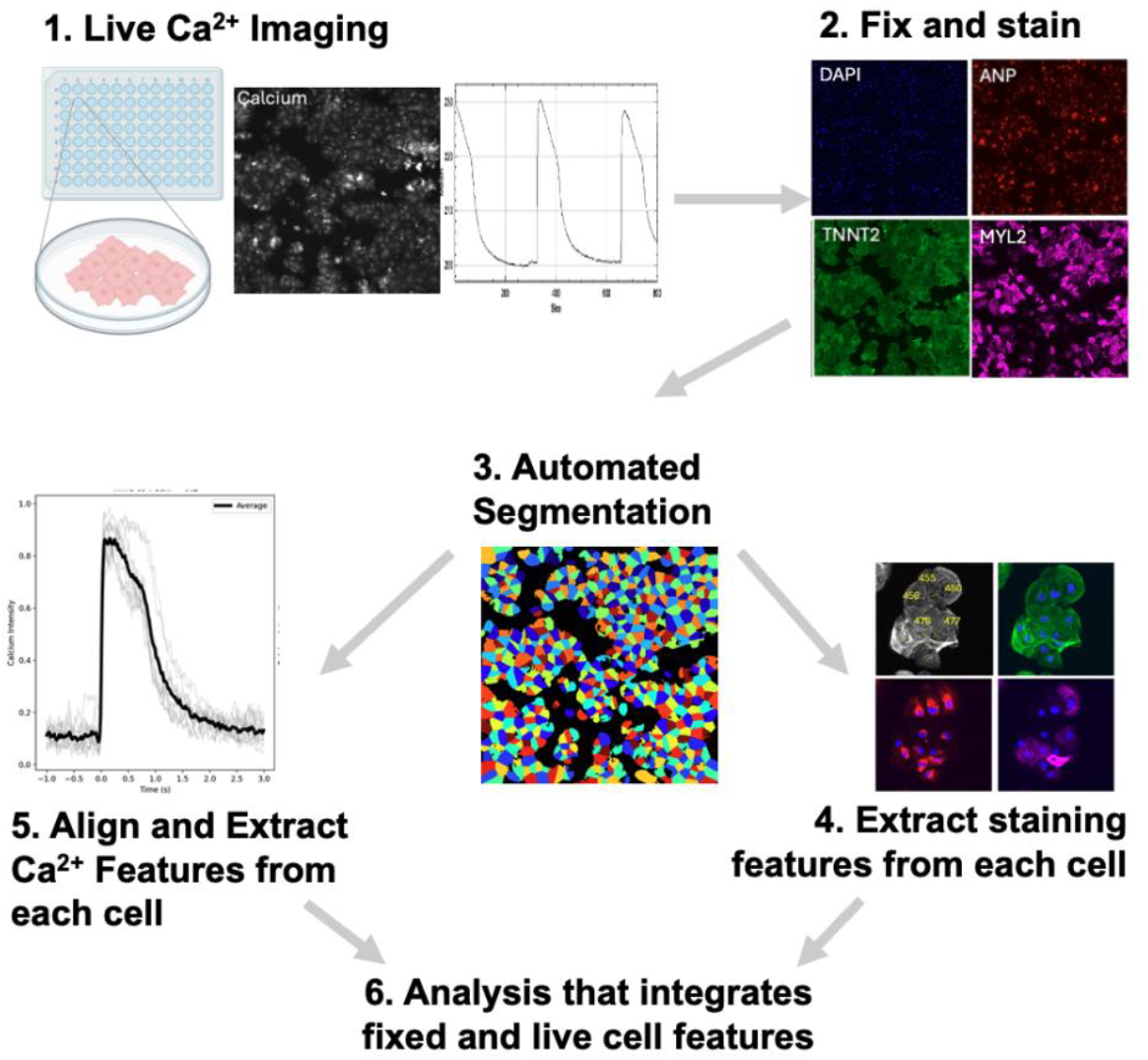
CARBONITE pipeline for quantitative single-cell analysis of live calcium and fixed staining images. In this study, iPSC-CMs are plated on a 96-well plate at a density of 20k cells / well. 1) Calcium transients are collected from each well, and then cells are fixed immediately after acquisition. 2) Cells are stained for DAPI (nuclear marker), TNNT2 (cardiomyocyte marker), ANP (perinuclear expression is atrial marker), and MYL2 (ventricular marker). **3)** An automated segmentation and imaging quantification tool called CellProfiler is used to segment the TNNT2. **4)** CellProfiler then quantifies single-cell features, including shape, and the relative expression and texture for DAPI, TNNT2, ANP, and MYL2 at the single-cell level. **5)** The TNNT2 images and segmentation are aligned to the Calcium transient video acquired in step 1, and single-cell calcium transients and transient features (e.g., calcium transient duration) extracted. **6)** Live (calcium) and fixed (staining) cell features are then investigated to understand the phenotypic heterogeneity and correlations across this multimodal dataset.

Experiments are performed in 96-well plates using a high-content automated imaging system (Molecular Devices ImageXpress) capable of rapid movement between wells. **1)** Live-cell calcium imaging is conducted at 37 °C using the calcium-sensitive dye Rhod-2, with short (10 s) recordings acquired per well to ensure consistent temporal sampling across the plate while minimizing phototoxicity. **2)** Immediately following calcium acquisition, cells are fixed and stained for nuclear DNA (DAPI) and protein markers defining cardiomyocyte identity, including the pan-cardiomyocyte marker TNNT2, the atrial marker ANP (with focus on perinuclear localization), and the ventricular marker MYL2.

**3)** Single-cell segmentation is performed using CellProfiler [17], with TNNT2 staining used to define cardiomyocyte boundaries. **4)** These segmentation masks are subsequently applied to extract per-cell shape features as well as quantitative measurements of protein expression intensity and spatial distribution. **5)** To enable accurate mapping between functional and molecular data, calcium videos are spatially aligned to post-fixation images using nuclear positions as anchors. This alignment step corrects for plate shifts of up to ∼30 μm that can occur during loading, ensuring reliable correspondence between calcium signals and stained cells. Following alignment, per-cell calcium traces are extracted and analyzed to quantify key dynamic features, including transient duration, time to peak, and decay kinetics.

**6)** These functional measurements are then statistically integrated with single-cell expression and shape metrics, enabling direct comparison of calcium dynamics across cardiomyocytes. While demonstrated here in a 96-well high-content imaging format, CARBONITE is broadly compatible with alternative plating configurations and microscopy platforms, providing a generalizable framework for paired functional and molecular single-cell phenotyping.

### Single-cell iPSC-CM features exhibit heterogeneity in calcium transient dynamics and marker expression

We applied CARBONITE to human iPSC-CMs known to have a mixed phenotype, extracting paired functional and molecular features at single-cell resolution (**Figure 2**). From each well, we quantified calcium transient properties alongside protein expression metrics for TNNT2, ANP, and MYL2 in the same cells, allowing direct assessment of cell-to-cell variability.

**Figure 2:**
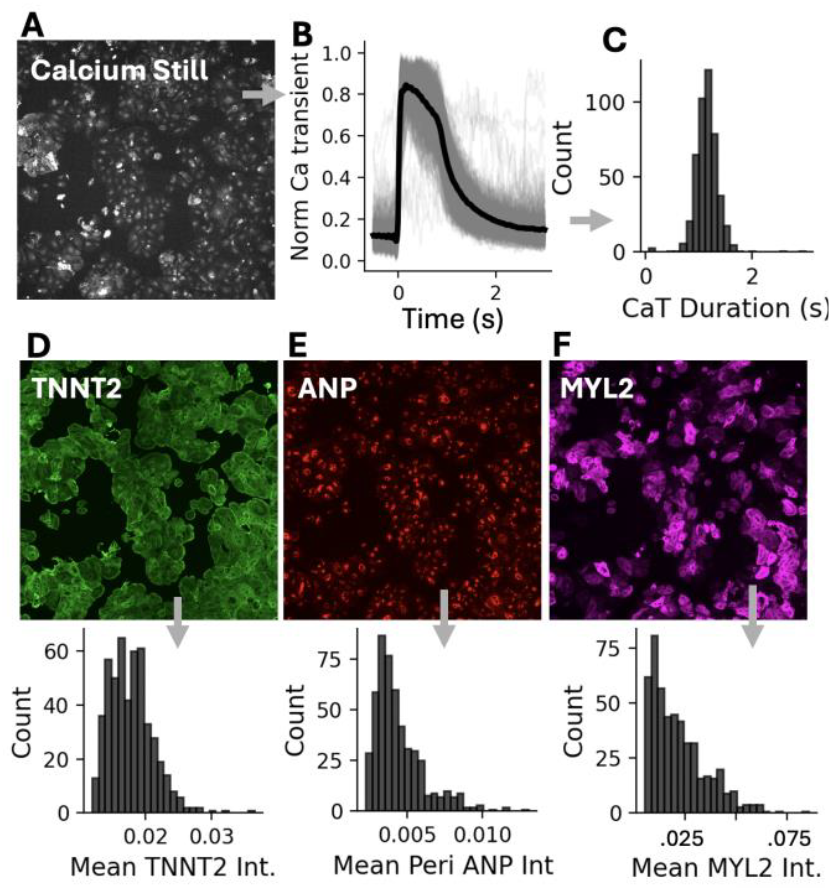
Extracting single-cell quantitative calcium transient and staining features from a single well. **A)** Still of the calcium video. **B)** Extracted single-cell calcium transients from video. **C)** Histogram of calcium transient duration (90% recovery to baseline) for all cells with calcium transients. **D-F)** TNNT2, ANP, and MYL2 images (top) and extracted mean cellular or perinuclear (for ANP) per-cell intensities.

**Figure 2** shows representative images from a single well, including a still frame from the calcium imaging video and the corresponding post-fixation immunostaining for TNNT2, ANP, and MYL2. Using CellProfiler-based segmentation, we computed per-cell calcium transient features and expression intensities, and visualized their distributions across the population. Histograms of calcium transient (CaT) duration (**Figure 2C**), mean perinuclear ANP intensity (**Figure 2E**), and mean MYL2 intensity (**Figure 2F**) reveal substantial heterogeneity among cells within the same well.

Notably, CaT duration and TNNT2 expression vaired widely but exhibited unimodal, near-normal distributions. In contrast, ANP and MYL2 expression displayed skewed distributions with extended right tails, reflecting smaller subpopulations of cells with markedly elevated atrial- or ventricular-associated marker expression. These long-tailed distributions suggest that cardiomyocyte subtype-associated molecular features are not uniformly expressed.

Together, these data demonstrate that single-cell calcium dynamics and identity marker expression vary widely within individual wells, motivating downstream analyses linking functional heterogeneity to molecular identity and spatial context.

### Canonical atrial and ventricular markers are associated with cell density and calcium transient shape

We next asked whether canonical atrial and ventricular cardiomyocyte markers—perinuclear ANP and MYL2, respectively—segregated cells into distinct groups and whether these markers were associated with other fixed shape features and live functional properties. Establishing such associations would support the use of CARBONITE to link molecular identity to emergent cellular behaviors, including calcium handling dynamics, at single-cell resolution.

Plotting perinuclear ANP intensity against MYL2 intensity revealed a negative correlation between these markers (**Figure 3A**), with cells distributed along an axis spanning high ANP/low MYL2 and low ANP/high MYL2 extremes. In addition to these two enriched populations, a substantial fraction of cells exhibited low expression of both markers. To specifically assess whether canonical atrial- and ventricular-associated markers corresponded to differences in other cellular features, we focused subsequent analyses on the two extreme groups (MYL2+/ANP- and MYL2-/ANP+, **Figure 3A**).

**Figure 3:**
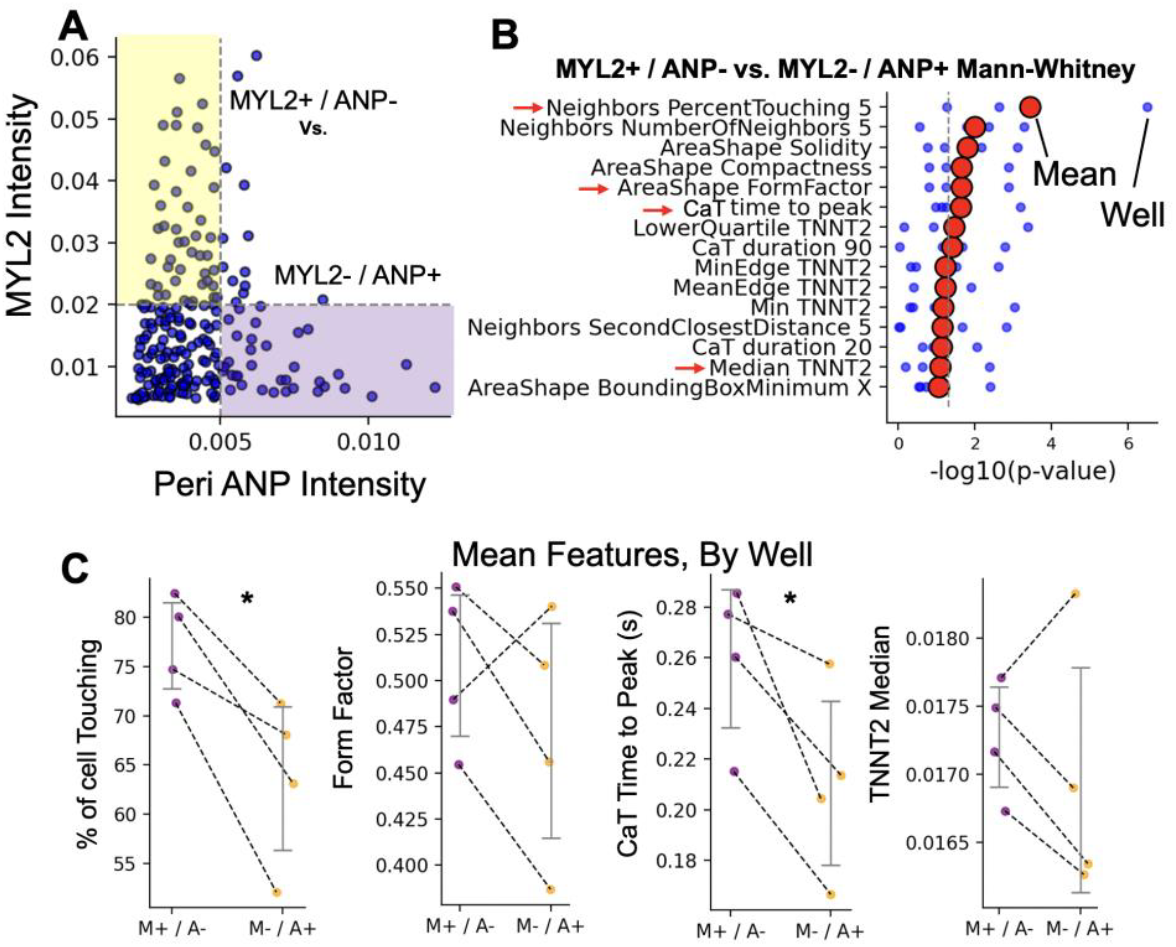
Features distinguishing MYL2+ / ANP- and MYL2- / ANP+ cardiomyocytes. **A)** Per-cell mean perinuclear ANP intensity vs. mean cellular MYL2 intensity. **B)** Per-well (Blue, n=4) and average (Red) p-values (Mann-Whitney U test) comparing MYL2+/ANP- (yellow in A) and MYL2-/ANP+ (purple in A) cardiomyocytes. **C)** Paired t-test for within-well average percent of cell touching adjacent cells (p=0.02), form factor (p=0.3), CaT time to peak (p=0.03), and TNNT2 median intensity (p=0.4).

For each well, we compared all quantified single-cell features between these two groups using a Mann– Whitney U test, yielding a p-value per feature per well (**Figure 3B**, blue points). We then computed the average p-value (red points) for each feature across wells, ranked features by −log10(p-value), and excluded ANP- and MYL2-derived measurements from downstream interpretation, as these were expected to strongly distinguish the groups by definition.

Among the top-ranked features was Neighbors Percent Touching 5, a CellProfiler-derived metric that quantifies the fraction of a cell’s boundary that is in direct physical contact with neighboring cells within a defined distance (here, five pixels). This measure reflects local cellular packing and contact-mediated neighborhood structure, indicating that cells enriched for atrial versus ventricular markers reside in distinct local contact environments. Specifically, ventricular-like (MYL2+/ANP-) cells tended to reside in more densely packed regions compared to atrial-like cells.

Additional features showing strong or moderate significance included select cell shape descriptors, calcium transient kinetics—most notably time to peak— and, to a lesser extent, TNNT2 intensity. To directly test these relationships, we performed paired t-tests across wells for four features spanning neighborhood context, cell shape, CaT dynamics, and TNNT2 expression (**Figure 3C**).

Consistent with the feature-ranking analysis, MYL2+/ANP-cells exhibited significantly higher Neighbors Percent Touching 5, indicating increased local cell density. In contrast, FormFactor, a shape metric that reflects how closely a cell approximates a perfect circle, did not differ significantly between groups, suggesting that gross cell shape was not a major determinant of marker-defined identity.

Calcium transient time to peak was significantly prolonged in MYL2+/ANP-cells, consistent with known differences in calcium handling kinetics between iPSC-CM derived atrial and ventricular cardiomyocytes [13]. Finally, median TNNT2 intensity did not differ significantly between groups, indicating that observed differences were not driven by variation in overall cardiomyocyte differentiation status.

Together, these analyses demonstrate that canonical atrial and ventricular markers are associated with distinct neighborhood contexts and calcium transient dynamics, while remaining largely independent of shape and cardiac troponin marker expression. These results support a model in which cardiomyocyte identity is coupled to local cellular organization and calcium handling phenotype, motivating subsequent analyses of spatial niches linking identity and function.

### Calcium transient shape defines two functional states associated with nuclear organization and ANP expression, but not MYL2

We next examined whether calcium transient dynamics were associated with cardiomyocyte molecular identity. Focusing on time to peak, a feature with significant difference between MYL2+/ANP- and MYL2-/ANP+ groups, we observed a bimodal distribution across single cells (**Figure 4A**). Cells segregated into two dominant populations characterized by fast (<0.2 s) or slow (>0.33 s) time to peak (**Figure 4B**), suggesting the presence of discrete functional states rather than a continuum. Visualization of calcium traces aligned to histogram-defined these groups revealed distinct transient morphologies. Cells in the fast group exhibited a sharp, spike-like rise followed by decay, whereas cells in the slow group displayed a broader, dome-shaped waveform with prolonged rise and decay phases (Figure 4B). These differences motivated a comparison across molecular and CaT shape features.

**Figure 4:**
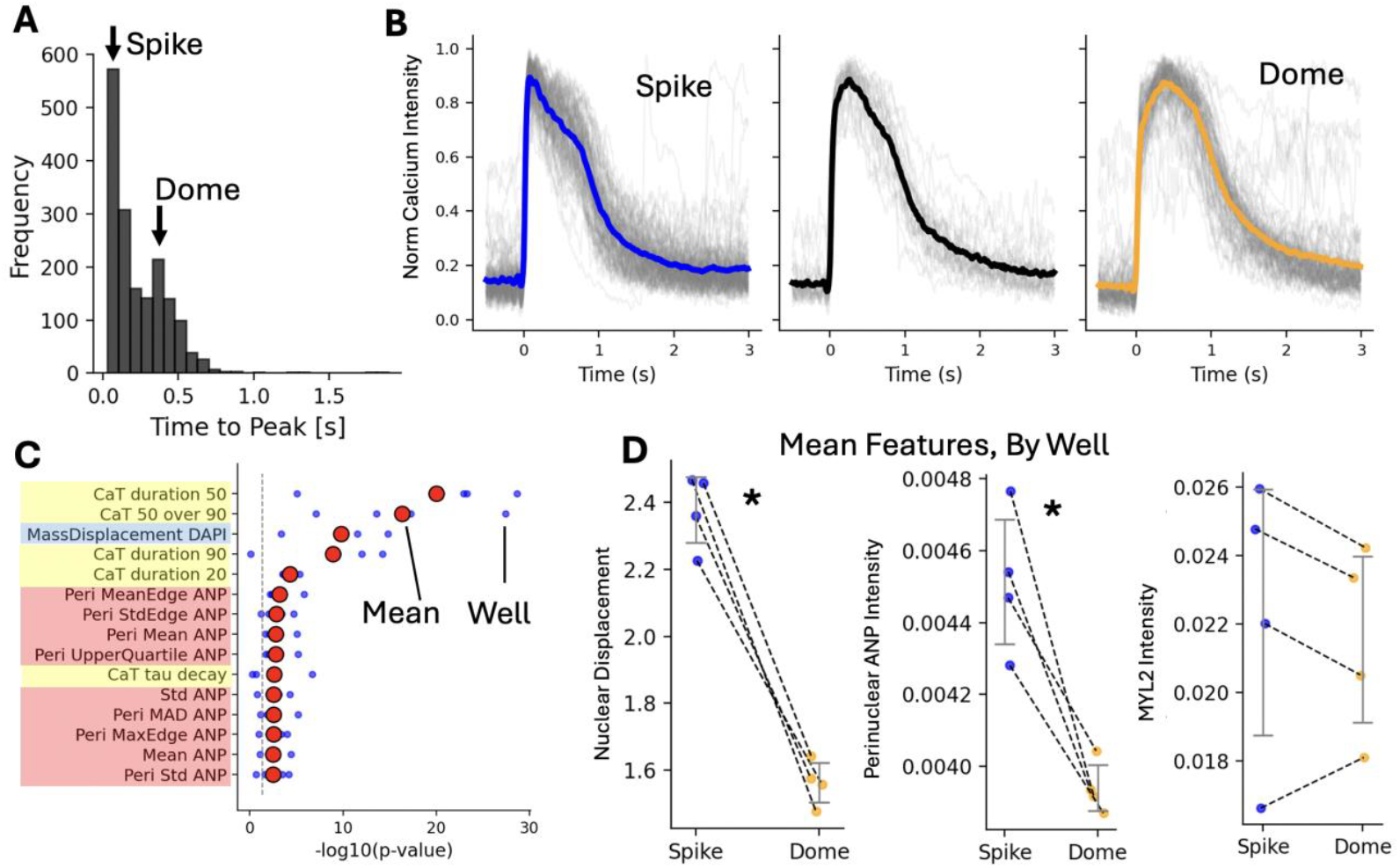
Nuclear mass displacement and perinuclear ANP intensity distinguish spike vs. dome CaT shape. **A)** Distribution of CaT time to peak for individual well shows bimodal distribution. **B)** CaT time to peak separated into three groups with the fast and slow groups showing distinct features: time to peak < 0.2 s (spike), time to peak > 0.33 s (dome), time to peak >0.2 s and <0.33 s (intermediate). **C)** Mann-Whitney U test identifies features that are significantly different in the spike vs dome group of cardiomyocytes – CaT features are highlighted in yellow, nuclear measures in blue, and ANP measures in red. Blue dots are within-well p-values and red dots are the average of four within-well p values. **D)** Paired t-test comparing the mean of nuclear displacement (p=0.001), mean perinuclear ANP intensity (p=0.01), and mean cellular MYL2 intensity (p=0.4).

To identify features distinguishing these two groups, we performed a Mann– Whitney U test across all quantified single-cell features (**Figure 4C**). As expected, multiple CaT-derived parameters ranked highly, reflecting intrinsic correlations among calcium dynamics metrics. Unexpectedly, however, several of the most strongly associated non-CaT features related to perinuclear ANP expression and nuclear spatial organization. In contrast, MYL2 intensity showed no significant association with time to peak, indicating that ventricular marker expression alone does not explain these functional differences.

Among the strongest non-CaT associations was nuclear mass displacement, defined as the distance between the cell centroid and the nuclear center of mass (computed from DAPI staining). Cells with spike-like CaT shape exhibited significantly greater nuclear mass displacement than cells with dome-shaped transients (paired t-test, **Figure 4D**). These spike-like cells also showed significantly higher perinuclear ANP enrichment, linking CaT morphology to atrial-associated expression patterns.

Closer inspection revealed that elevated nuclear mass displacement primarily reflected binucleation, rather than asymmetric positioning of a single nucleus. Because the CellProfiler pipeline assumes a one-to-one correspondence between cell and nucleus, closely spaced nuclei within binucleated cells lead to apparent displacement of the nuclear center of mass relative to the cell centroid. Cells with the highest nuclear displacement were predominantly binucleated. To more directly quantify this property, we measured inter-nuclear distance for all cells – binucleated cells are expected to have a smaller inter-nuclear distance than adjacent mononucleated (**Figure 5**). This is what we see – a smaller inter-nuclear distance for the spike-shaped CaT cells.

**Figure 5:**
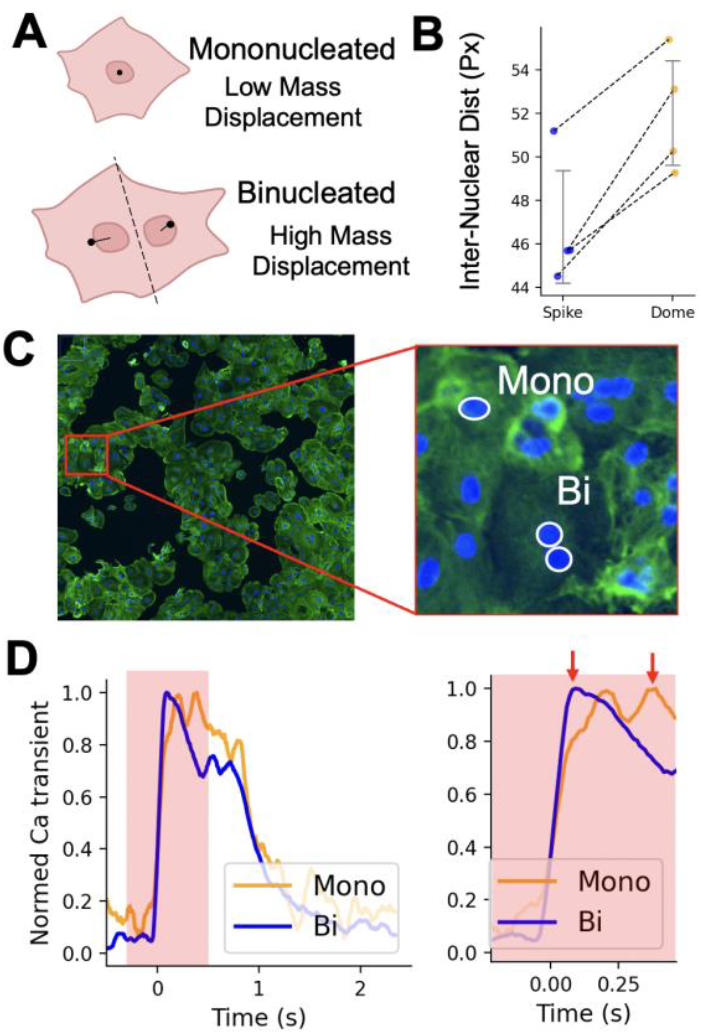
Nucleation state distinguishes spike vs. dome CaT phenotypes. **A)** Centroids (black dot) in mononucleated cells are near center of nucleus. CellProfiler segments binucleated cells into two cells, creating a border between the two nuclei – this creates an increased distance between the cellular centroid and nuclear center of mass. **B)** Inter-nuclear distance is significantly less (p=0.009) in spike vs. dome CaT. **C)** TNNT2+DAPI image (left) and zoom (right) shows adjacent mono and binucleated cardiomyocytes **D)** CaT (left) and zoomed CaT rise phase (right) for the mononcleated and binucleated iPSC-CMs in C. Red arrows (right) denote the peaks for mono and binucleated CMs.

Together, these results demonstrate that calcium transient morphology segregates cardiomyocytes into two discrete functional states that are strongly associated with nucleation state and perinuclear ANP expression, but surprisingly not with MYL2 expression. This dissociation between ventricular marker expression and calcium dynamics highlights the importance of single-cell, paired functional–molecular measurements for resolving the multiple dimensions of cardiomyocyte identity that contribute to heterogeneity.

## Discussion

In this study, we introduce CARBONITE, a scalable framework that directly links live calcium dynamics with molecular and spatial features in the same individual cardiomyocytes. Applying this approach to mixed iPSC-CM populations reveals that functional heterogeneity in calcium handling is highly structured rather than random. Calcium transient dynamics and molecular marker expression vary widely within individual wells, yet this variability follows reproducible patterns across cells and experiments. By resolving these patterns at single-cell resolution, CARBONITE identifies functional states that are obscured by population-averaged analyses. Together, these findings establish CARBONITE as a platform for identifying biologically meaningful organization within heterogeneous cardiomyocyte populations and motivate deeper interrogation of the axes linking function, identity, and cellular context.

### Atrial–ventricular identity: markers and electrophysiology are informative but incomplete

Canonical atrial and ventricular markers [6], together with electrophysiological features such as action potential (AP) shape [14], are widely used to stratify cardiomyocytes. Implicit in these approaches is the assumption that molecular subtype, AP morphology, and underlying functional state are tightly coupled. Additionally, the iPSC-CM electrophysiological computational models assume that AP feature-based grouping is sufficient to build chamber-specific models [18]. More recent models incorporate heterogeneity, but do not explain this heterogeneity with underlying molecular identity [19]. Our results support this framework in part, as ANP and MYL2 expression associate with some differences in calcium handling and local cellular context. However, these features capture only a subset of functional variability. Notably, ANP enrichment correlates strongly with calcium transient shape, whereas MYL2 expression does not explain calcium dynamics. More broadly, prior single-cell electrophysiology studies have shown that AP shape reflects the combined influence of multiple ionic currents rather than a single molecular identity axis [8]. Together, these findings indicate that marker- and AP-based classifications represent projections of a higher-dimensional identity space, and that reliance on either alone may obscure additional axes of cardiomyocyte function.

### Nucleation as a dominant, previously underappreciated axis of calcium function

Beyond canonical subtype markers, we identify nucleation state as a major determinant of calcium transient shape. Cells exhibiting spike-like calcium transients are strongly enriched for multinucleation, whereas dome-shaped transients are predominantly observed in mononucleated cells. While multinucleation is a well-known feature of cardiomyocytes, its functional consequences have been difficult to quantify at single-cell resolution. Previous work has shown electrophysiological and functional differences based on nucleation status in cardiomyocytes, but this work does not compare function across chamber/subtypes [20,21]. Our results demonstrate that nucleation state explains calcium transient shape more effectively than ventricular marker expression, revealing an unexpected axis of functional identity. These findings suggest that cellular organization and structural state—rather than transcriptional subtype alone—play a central role in shaping calcium handling dynamics. Nucleation thus represents a previously underappreciated contributor to cardiomyocyte functional heterogeneity that warrants further mechanistic investigation.

### CARBONITE is a tool for functional multimodal single-cell biology

CARBONITE conceptually parallels multimodal single-cell genomics by integrating multiple orthogonal measurements within the same cell, but extends this paradigm to dynamic physiological function. Whereas single-cell RNA-seq and ATAC-seq resolve transcriptional and epigenetic heterogeneity, CARBONITE captures quantitative calcium dynamics alongside molecular identity and spatial context. This integration enables direct mapping between functional states and molecular features, rather than inference across separate cell populations. Importantly, CARBONITE achieves this without genetic reporters, specialized substrates, or destructive assays, making it broadly compatible with standard imaging platforms and culture formats. As a result, CARBONITE provides a generalizable framework for functional multimodal single-cell analysis, enabling discovery of organizational principles governing cardiomyocyte behavior that are inaccessible to single-modality approaches.

### Limitations and future directions

Several limitations should be considered when interpreting these findings. First, CARBONITE currently focuses on calcium dynamics and a limited panel of protein markers, and does not capture membrane voltage or broader transcriptional state. Second, measurements were performed in two-dimensional cultures and over short recording windows. These constraints, however, are largely technical rather than conceptual. Future extensions could incorporate voltage [12] imaging or expanded immunophenotyping [22,23]. Additionally, applying CARBONITE under genetic or pharmacological perturbations will enable causal interrogation of the axes identified here. CARBONITE provides a scalable and adaptable foundation for dissecting cardiomyocyte identity across functional, molecular, and organizational dimensions.

## Acknowledgements

This work was supported by the American Heart Association with a grant (25POST1356494) to A.P.C. and the National Heart Lung and Blood Institutes with grants R01HL160665 and R01HL162925 to J.J.S.

## Materials and Methods

### Human iPSC-CM culture

Human iPSC-derived cardiomyocytes (iCell® Cardiomyocytes_^2^_; Fujifilm Cellular Dynamics, Inc.) were thawed according to the manufacturer’s instructions and plated into 96-well plates at a density of 15,000 cells per well using iCell plating medium. Cells were maintained in iCell maintenance medium, with complete media exchanges performed every two days. All cultures were maintained under standard incubator conditions (37 °C, 5% CO2). Calcium imaging experiments were initiated one week after thaw to allow recovery and stabilization of spontaneous beating activity.

### Calcium imaging acquisition

For calcium imaging, cells were first washed once with Hanks’ Balanced Salt Solution (HBSS) and then loaded with the calcium-sensitive dye Rhod-2 AM (Thermo Fisher Scientific) at a final concentration of 4 μM diluted in Tyrode’s solution. Cells were incubated with Rhod-2 at 37 °C for 25 minutes to allow dye loading. Following incubation, cells were washed with Tyrode’s solution and returned to fresh Tyrode’s for an additional 5–10 minutes in the incubator to permit recovery prior to imaging. All calcium imaging was performed in Tyrode’s solution under physiological temperature (35-37º C) conditions. Images were collected using a Molecular Devices ImageXpress with magnification set to 10x and at a frame rate of 100Hz. Calcium was collected for 10 s per well to allow the camera to image multiple wells on the plate.

### Fixation and immunofluorescence staining

Immediately following calcium imaging, cells were fixed and processed for immunofluorescence staining using a one-day protocol. Cells were permeabilized in PBS containing 0.2% Triton X-100, followed by thorough rinsing with Dulbecco’s PBS (D-PBS). Cells were then blocked in PBS containing 1% bovine serum albumin (BSA) to reduce nonspecific binding. Primary antibodies against TNNT2, ANP, and MYL2 were diluted in blocking solution and applied to cells. After primary antibody incubation, cells were washed with PBS and incubated with species-specific donkey secondary antibodies conjugated to Alexa Fluor dyes. Nuclei were counterstained with DAPI prior to final washes and imaging. All staining steps were performed directly in 96-well plates. Images were collected using the Molecular Devices ImageXpress.

### Image segmentation and feature extraction

Single-cell image segmentation and feature extraction were performed using CellProfiler. Nuclei were first identified from DAPI staining and used as seeds to define individual cells. Cardiomyocyte boundaries were then delineated by propagating outward from each nucleus using TNNT2 staining to generate cell masks. A perinuclear region was defined as a fixed-width annulus extending six pixels from the nuclear boundary. Using these segmentation masks, per-cell measurements were extracted for TNNT2, ANP, and MYL2 intensity, as well as perinuclear ANP enrichment and nuclear (DAPI) features. In addition to intensity-based metrics, CellProfiler was used to compute cell shape and neighborhood features for downstream analyses.

### Calcium transient extraction and analysis

To enable per-cell calcium transient quantification, live calcium imaging videos were manually aligned to post-fixation DAPI and TNNT2 images to correct for small spatial shifts introduced during plate handling. Nuclear centroids identified from DAPI segmentation were used as reference points for calcium signal extraction. For each cell, calcium intensity was quantified by averaging fluorescence within a 7 × 7 pixel region centered on the nuclear centroid for each frame of the calcium imaging video, generating a single calcium transient trace per cell. Traces were normalized to a 0–1 range and temporally smoothed to reduce high-frequency noise using frequency-domain low-pass filtering followed by moving-average smoothing. From the processed calcium transients, dynamic features including time to peak, transient duration, and decay kinetics were extracted for downstream statistical analyses. This approach enabled robust, nucleus-referenced quantification of calcium dynamics at single-cell resolution while minimizing contamination from neighboring cells.

### Statistical analysis

All statistical analyses were performed at the single-cell level with well-level replication explicitly preserved. For analyses comparing calcium transient morphologies, cells were stratified into two functional groups based on calcium transient time to peak: fast, spike-like transients (<0.2 s) and slow, dome-shaped transients (>0.33 s). Within each well, distributions of quantitative cellular features were compared between groups using two-sided Mann–Whitney U tests, generating a p-value per feature per well. Features were then ranked based on the mean −log10(p-value) across wells to identify reproducible associations while minimizing the influence of any single well. This same approach was used when comparing the MYL2+ / ANP- and MYL2- / ANP+ groups.

For selected features of interest, including molecular, spatial, and nuclear organization metrics, well-level means were computed separately for spike- and dome-shaped cells. These paired measurements were compared across wells using paired t-tests, enabling direct assessment of consistent within-well differences between functional states. Statistical significance was assessed using two-sided tests, and results are reported as mean ± standard deviation unless otherwise indicated. This same approach was used when comparing the MYL2+ / ANP- and MYL2- / ANP+ groups.

## References

1. Clark AP, Wei S, Kalola D, Krogh-Madsen T, Christini DJ. An in silico-in vitro pipeline for drug cardiotoxicity screening identifies ionic pro-arrhythmia mechanisms. Br J Pharmacol. 2022;179:4829–4843. doi:10.1111/bph.15915

2. Zhu L, Choudhary K, Gonzalez-Teran B, Ang Y-S, Thomas R, Stone NR, et al. Transcription Factor GATA4 Regulates Cell Type-Specific Splicing Through Direct Interaction With RNA in Human Induced Pluripotent Stem Cell-Derived Cardiac Progenitors. Circulation. 2022;146:770–787. doi:10.1161/CIRCULATIONAHA.121.057620

3. Goversen B, Jonsson MK, van den Heuvel NH, Rijken R, Vos MA, van Veen TA, et al. The influence of hERG1a and hERG1b isoforms on drug safety screening in iPSC-CMs. Prog Biophys Mol Biol. 2019;149:86–98. doi:10.1016/j.pbiomolbio.2019.02.003

4. Schocken D, Stohlman J, Vicente J, Chan D, Patel D, Matta MK, et al. Comparative analysis of media effects on human induced pluripotent stem cell-derived cardiomyocytes in proarrhythmia risk assessment. J Pharmacol Toxicol Methods. 2018;90:39–47. doi:10.1016/j.vascn.2017.11.002

5. Blinova K, Stohlman J, Vicente J, Chan D, Johannesen L, Hortigon-Vinagre MP, et al. Comprehensive Translational Assessment of Human-Induced Pluripotent Stem Cell Derived Cardiomyocytes for Evaluating Drug-Induced Arrhythmias. Toxicol Sci. 2017;155:234–247. doi:10.1093/toxsci/kfw200

6. Kathiriya IS, Rao KS, Iacono G, Devine WP, Blair AP, Hota SK, et al. Modeling Human TBX5 Haploinsufficiency Predicts Regulatory Networks for Congenital Heart Disease. Developmental Cell. 2021;56:292-309.e9. doi:10.1016/j.devcel.2020.11.020

7. Kathiriya IS, Rao KS, Clark AP, Hu KM, Grant ZL, Matthews MN, et al. Reduced TBX5 dosage undermines developmental control of atrial cardiomyocyte identity in a model of human atrial disease. Development. 2026;153:dev205173. doi:10.1242/dev.205173

8. Clark AP, Wei S, Fullerton K, Krogh-Madsen T, Christini DJ. Single-cell ionic current phenotyping explains stem cell-derived cardiomyocyte action potential morphology. Am J Physiol Heart Circ Physiol. 2024;326:H1146–H1154. doi:10.1152/ajpheart.00063.2024

9. Clark AP, Krogh-Madsen T, Christini DJ. Stem cell-derived cardiomyocyte heterogeneity confounds electrophysiological insights. J Physiol. 2024. doi:10.1113/JP284618

10. Clark AP, Clerx M, Wei S, Lei CL, de Boer TP, Mirams GR, et al. Leak current, even with gigaohm seals, can cause misinterpretation of stem cell-derived cardiomyocyte action potential recordings. Europace. 2023;25:euad243. doi:10.1093/europace/euad243

11. Feyen DAM, McKeithan WL, Bruyneel AAN, Spiering S, Hörmann L, Ulmer B, et al. Metabolic Maturation Media Improve Physiological Function of Human iPSC-Derived Cardiomyocytes. Cell Rep. 2020;32:107925. doi:10.1016/j.celrep.2020.107925

12. Klimas A, Ambrosi CM, Yu J, Williams JC, Bien H, Entcheva E. OptoDyCE as an automated system for high-throughput all-optical dynamic cardiac electrophysiology. Nat Commun. 2016;7:11542. doi:10.1038/ncomms11542

13. Cyganek L, Tiburcy M, Sekeres K, Gerstenberg K, Bohnenberger H, Lenz C, et al. Deep phenotyping of human induced pluripotent stem cell-derived atrial and ventricular cardiomyocytes. JCI Insight. 2018;3:e99941. 99941. doi:10.1172/jci.insight.99941

14. Ma J, Guo L, Fiene SJ, Anson BD, Thomson JA, Kamp TJ, et al. High purity human-induced pluripotent stem cell-derived cardiomyocytes: electrophysiological properties of action potentials and ionic currents. Am J Physiol Heart Circ Physiol. 2011;301:H2006–2017. doi:10.1152/ajpheart.00694.2011

15. Churko JM, Garg P, Treutlein B, Venkatasubramanian M, Wu H, Lee J, et al. Defining human cardiac transcription factor hierarchies using integrated single-cell heterogeneity analysis. Nat Commun. 2018;9:4906. doi:10.1038/s41467-018-07333-4

16. Muncie-Vasic JM, Sinha T, Clark AP, Brower EF, Saucerman JJ, Black BL, et al. MEF2C controls segment-specific gene regulatory networks that direct heart tube morphogenesis. Genes Dev. 2025. doi:10.1101/gad.352889.125

17. Stirling DR, Swain-Bowden MJ, Lucas AM, Carpenter AE, Cimini BA, Goodman A. CellProfiler 4: improvements in speed, utility and usability. BMC Bioinformatics. 2021;22:433. doi:10.1186/s12859-021-04344-9

18. Paci M, Hyttinen J, Aalto-Setälä K, Severi S. Computational models of ventricular- and atrial-like human induced pluripotent stem cell derived cardiomyocytes. Ann Biomed Eng. 2013;41:2334–2348. doi:10.1007/s10439-013-0833-3

19. Kernik DC, Morotti S, Wu H, Garg P, Duff HJ, Kurokawa J, et al. A computational model of induced pluripotent stem-cell derived cardiomyocytes incorporating experimental variability from multiple data sources. J Physiol. 2019;597:4533–4564. doi:10.1113/JP277724

20. Huang C-F, Chen Y-C, Yeh H-I, Chen S-A. Mononucleated and binucleated cardiomyocytes in left atrium and pulmonary vein have different electrical activity and calcium dynamics. Prog Biophys Mol Biol. 2012;108:64–73. doi:10.1016/j.pbiomolbio.2011.09.007

21. Kiessling P, Joodaki M, Pikulska D, Scheidereit E, Cesaro G, Ruiz M, et al. Polyploid cardiomyocytes define disease-specific transcriptional states in the mammalian heart. 2026. doi:10.64898/2026.01.31.701472

22. Chang Q, Ornatsky OI, Siddiqui I, Loboda A, Baranov VI, Hedley DW. Imaging Mass Cytometry. Cytometry A. 2017;91:160–169. doi:10.1002/cyto.a.23053

23. Baharlou H, Canete NP, Cunningham AL, Harman AN, Patrick E. Mass Cytometry Imaging for the Study of Human Diseases-Applications and Data Analysis Strategies. Front Immunol. 2019;10:2657. doi:10.3389/fimmu.2019.02657

